# DLPAlign: A Deep Learning based Progressive Alignment for Multiple Protein Sequences

**DOI:** 10.1101/2020.07.16.207951

**Authors:** Mengmeng Kuang

## Abstract

This paper proposed a novel and straightforward approach to improve the accuracy of progressive multiple protein sequence alignment. We trained a decision-making model based on the convolutional neural networks and bi-directional long short term memory networks, and based on this model, we progressively aligned the input sequences by calculating different posterior probability matrixes.

To test the accuracy of this approach, we have implemented a multiple sequence alignment tool called DLPAlign and compared its performance with eleven leading alignment methods on three empirical alignment benchmarks (BAliBASE, OXBench and SABMark). Our results show that DLPAlign can get the best total-column scores on the three benchmarks. When evaluated against the 711 low similarity families with average PID ≤ 30%, DLPAlign improved about 2.8% over the second-best MSA software.

Besides, we also compared the performance of DLPAlign and other alignment tools on a real-life application, namely protein secondary structure prediction on four protein sequences related to SARS-COV-2, and DLPAlign provides the best result in all cases.

## I. Introduction

A Multiple Sequence Alignment (MSA) can be seen as a table constructed from protein sequences with an appropriate amount of spaces inserted. MSA can be applied in many cases, such as recovering the history or relationship between protein or amino acid sequences and digging out some structural or functional roles of the sequences [1]. More and more biological modeling methods rely on the assembly of precise MSAs [2]. Therefore, it is of considerable significance to construct an algorithm to assist the MSA construction.

The MSA construction problem can be defined as a mathematical problem. Given *n* sequences *S_i_*, *i* = 1, 2, ⋯, *n* as Equation (1),

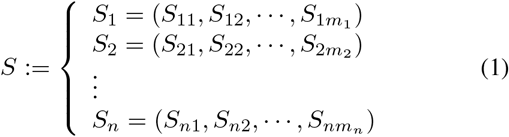

an MSA is constructed from this set of sequences by inserting an appropriate amount of gaps needed into each of the *S_i_* sequences of *S* until the modified sequences, 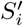, all conform to a same length *l* and no values in the sequences of *S* of the same column *m*, consists of only gaps. The mathematical form of an MSA of the above sequence set is shown at Equation (2):

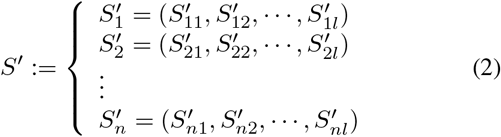

For a protein family, if the number of sequences is small and the length is short, we may be able to align it manually, but when we face a large number of sequences or long sequences, we need to use computer programs to handle it.

Since the early 1980s, the MSA construction problem has been solved by the algorithm-centric approach [2]: Design algorithms to solve the combinatorial optimization problem, which is to find the alignment with the most massive sum of column scores. Many excellent algorithms are well applied, such as dynamic programming [3], divide and conquer algorithm [4] and so on. Besides, in the past few decades, many alignment strategies have been proposed, such as progressive strategy [5], non-progressive strategy, consistency-based method [6], iterative refinement [7] etc.

The progressive strategy is one of the maturest MSA strategies with large amounts of research validation and highest accuracy. A typical MSA method using progressive strategy mainly includes five parts: (1) posterior probability matrix calculation, (2) distance matrix calculation, (3)“Guide Tree” [8] generation by clustering methods, (4) consistency transformation, and (5) refinement. In past studies, the main research focus is on the calculation method of the posterior probability matrix, the generation method of the guide tree, and the consistency transformation method.

The most popular MSA tools adopting a progressive strategy in the past 10 years are as follows: (1) ProbCons [9] uses a pair-hidden Markov model (HMM) to calculate the posterior probability matrix, an unweighted probabilistic consistency transformation, using an unweighted pair group method with the arithmetic mean (UPGMA) [10] hierarchical clustering method to generate a guide tree and iterative refinement to construct an MSA. (2) Probalign [11], another popular, highly accurate MSA tool, uses a partition function instead of the ProbCons pair HMM to calculate the posterior probability matrix. (3) MSAProbs [12] combines (1) and (2), using the Root Mean Square (RMS) of pair HMM and the partition function as the calculation method for the posterior probability matrix, and adopts a weighted consistency transformation. (4) GLProbs [13] introduced random HMM, and adaptively uses (i) the partition function, (ii) global pair HMM, (iii) the RMS of global pair HMM and random HMM to calculate the posterior probability matrix using the different average pairwise percentage identity (PID) of each protein family. PID stands for the percentage of the number of homologous positions in the pairwise alignment of two sequences. (5) PnpProbs [14] applies UPGMA and the weighted pair group method with arithmetical mean (WPGMA) [15] adaptively to generate the guide tree in its progressive branch.

Although these MSA tools can achieve relatively high accuracy on the whole, when it comes to a specific protein family, the accuracy of the different tools is often different [16]. Most importantly, after so many years of research, the algorithm-centric method has not significantly improved accuracy. As for the progressive alignment strategy, protein families with low similarity have always been the most challenging part [17].

This paper explores a different approach, namely the data-centric approach [18], to tackle the MSA construction problem to further improve the accuracy of MSAs, especially the accuracy on low similarity protein families. We first determined which specific part in the progressive MSA methods could provide the greatest improvement in accuracy. Then we applied deep learning methods to train decision-making models to choose the most appropriate calculation method for that part.

*Our contributions:* After determined which specific part in progressive MSA methods to improve, we transformed the classification of MSA families into the classification of sequence pairs, and in this process, we obtained large-scale training data (there were 954854 such training data). We gained these protein sequences data from many datasets which were SISYPHUS [19], SABmark [20], BAliBASE [21] and a part of the extension set of BAliBASE namely BAliBASE-X, OXBench [22], HOMSTRAD [23], and Mattbench [24]. Table I summarizes, for each dataset, the number of families and the total number of sequences.

**TABLE I:** The number of families and the total number of sequences in each dataset.

|  | Num. of families | Total num. of sequences |
| --- | --- | --- |
| BALiBASE | 386 | 11082 |
| BALiBASE-X | 147 | 6031 |
| OXBench | 395 | 3292 |
| SABmark | 423 | 2418 |
| SISYPHUS | 126 | 1772 |
| HOMSTRAD | 1030 | 3454 |
| Mattbench | 259 | 1698 |
| **Total** | **2766** | **29747** |

Deep learning methods were used to train decision-making models to help us choose the most appropriate calculation method of the specific part. We give more details of the decision-making models and the implementation in Sections II-B and III-B.

Further, based on the most accurate decision-making model, we build a new progressive MSA tool called DLPAlign. We compared DLPAlign with eleven popular MSA tools including PnpProbs, QuickProbs [25], GLProbs, PicXAA [26], Prob-Cons, MSAProbs, MAFFT [27], Muscle [28], ClustalΩ [29], ProbAlign and T-Coffee [30] on three empirical benchmarks SABmark, BAliBASE and OXBench. Tables V, VI and VII (in Section IV) demonstrate the alignment accuracies of the MSAs constructed by the tools for families in BAliBASE, OXBench and SABMark, respectively. DLPAlign achieved the highest TC-scores among all the tools on all benchmarks.

Note that DLPAlign got better performance on the low or medium similarity protein families that other progressive methods were not good at. Figures 1(a) and 1(b) compare the average TC-scores on low similarity (with PID ≤ 30% and there are totally 711 such families) and medium similarity (with 30% ¡ PID ≤ 60% and there are totally 352 such families) families in BAliBASE, OXBench and SABmark. It can be seen from the figures that the improvement of DLPAlign is pronounced, especially in the low similarity families set.

**Fig. 1:**
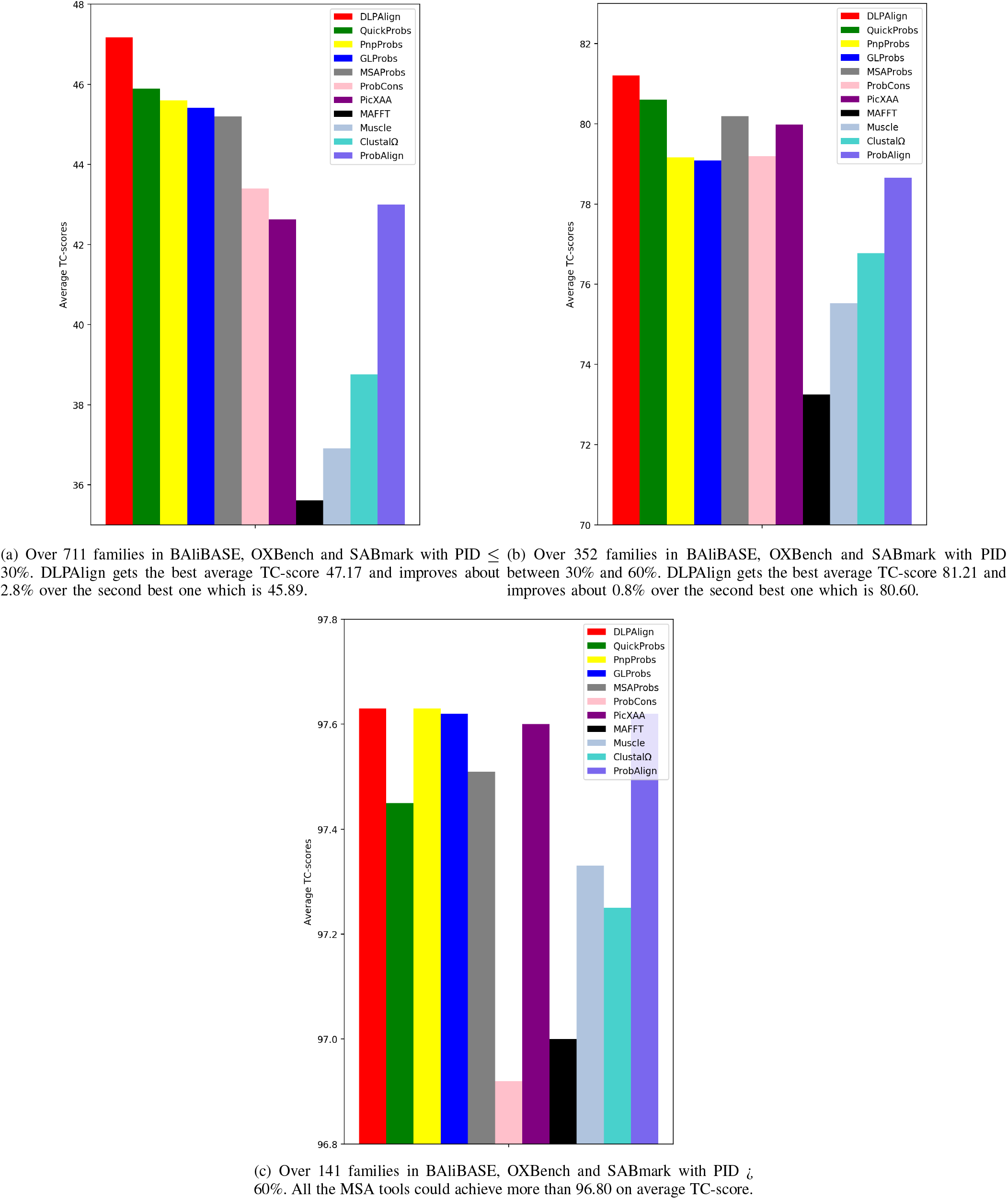
Comparison of average TC-scores over different similarity protein families in BAliBASE, OXBench and SABmark

We think this tool can be used in actual MSA task, so we upload the source code as well as the benchmarks for testing to GitHub (https://github.com/kuangmeng/DLPAlign).

## II. Methods

In this section, we first explain how to determine which part is used to improve the accuracy, then consider which data to use as the training data, and finally introduce the decisionmaking process of our deep learning methods.

### A. How to select the best promotion part?

As we mentioned in Section I, a typical progressive alignment method consists of five main parts: (1) posterior probability matrix calculation, (2) distance matrix calculation, (3) “Guide Tree” generation by clustering methods, (4) consistency transformation and (5) refinement. The most studied of these parts are the posterior probability matrix calculation, clustering methods for generating guide tree, and consistency transformation. We chose these three parts from Parts (1) - (5) as our candidate promotion parts and refer to the posterior probability matrix calculation as **Part A**, guide tree generation as **Part B**, and consistency transformation as **Part C**. For each part, we extracted several candidate options from previous studies [9], [11], [12], [13], [14], as shown below:

Options for Part A:

1. Pair-HMM
2. Partition function
3. the RMS of pair-HMM and partition function
4. the RMS of pair-HMM, partition function and random HMM

Options for Part B:

1. UPGMA
2. WPGMA

Options for Part C:

1. Unweighted consistency transformation
2. Weighted consistency transformation

When we test a single part, we need to use the same methods for the other parts. We used the calculation method of each part numbered (1) as the default method for that part. For other parts that we did not select, such as the distance matrix calculation and refinement, we adopted the same implementation method as in GLProbs.

The critical concern is the upper bounds of the various calculations in different parts of the progressive alignment strategy, and in which part the maximum improvement can be made. For the *i*-th option of Part A, we implemented a pipeline 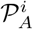. We obtained pipelines 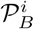 and 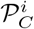 in the same way. we report the results of these pipelines in Section III-A.

### B. How to train decision-making models?

In **Part X** of the progressive alignment strategy, for a specific protein family, 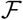, choosing the method with the highest accuracy on which to construct an MSA 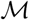 can be expressed as the classification problem 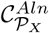. Classification 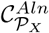 of a protein family is defined as follows:

> 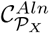has *n* classes 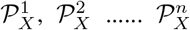. A protein family 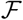 is in class 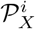 if the MSA constructed by the pipeline 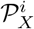 could get better TC-score than those constructed by others, where *n* is the number of options in Part X and *i* is a positive integer not greater than *n*.

#### 1) Data augmentation

In the past few years, there have been significant developments in deep learning, which have been applied in Bioinformatics [31], mainly due to the continuous expansion of the data scale. It is not sufficient to use 2766 protein families or 29747 sequences to train deeplearning models, so we considered coupling any two sequences in the same protein family as an independent piece of data. If a family has *n* sequences, we can get 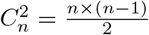 sequence pairs from it. In this way, our data was expanded to 954,854 pairs.

#### 2) The structure of candidate deep learning models

Because the length of the sequence pairs was not very consistent, we normalized the length before choosing the neural networks. We unified all pairs into a fixed length *α* (*α* = 512 in our structure). When the length of a pair was insufficient, it was filled by gaps at the end to increase the length to the *α*. When the length of the sequence exceeded *α*, only the leading fragments of length *α* were intercepted.

When we regard each character as a single word, if we convert it into a one-hot word vector, the size of the vector is a little large, so we first used the word-embedding [32] technique to convert each word into a small size (eight-dimensional vector).

Even so, our input scale was still relatively large, so we applied convolutional neural networks (CNNs) [33], which had made a significant breakthrough in computer vision to reduce the dimensionality of the data while retaining its characteristics, as the first two layers of our models.

There were order relationships between every character in a protein sequence pair, so we added a recurrent neural network (RNN) [34] layer after the CNNs. The improved versions of the recurrent neural network, long short term memory network (LSTM) [35] and gated recurrent unit network (GRU) [36], and their bi-directional versions (BiLSTM, BiGRU) have many advantages, so they were alternatives.

Subsequently, two full connection layers were connected. To reduce overfitting, we added a specific dropout rate to the first full connection layer.

This kind of neural network structure is very suitable and widely used for classification tasks [37], [38], [39], [40].

Section III-B gives the implementation of different deep learning models, as well as the training and testing processes. The final decision-making model was determined according to the accuracy of different models.

## III. Implementation

To test the effectiveness of our methods and select the best promotion part, we have implemented pipelines 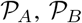 and 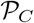 by adopting different calculation methods in Part A, Part B and Part C of progressive alignment strategy. In the meantime, different deep learning models were trained for choosing the best decision-making model for the best promotion part Part X. All these efforts were to implement DLPAlign.

### A. Find the best promotion part

To evaluate the advantages and disadvantages of several methods in Part A and to what extent they could be improved, we got four different pipelines by using different calculation methods of the posterior probability matrix in the GLProbs’ code and implemented the calculation in Parts B and C by default, naming them 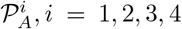, which respectively represents the different options of Part A mentioned above. We implemented 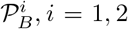, where *i* denoted the different clustering methods for guide tree generation in Part B, and 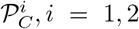, where *i* denoted the different calculation of consistency transformation in Part C in the same way.

To measure the accuracies of MSAs constructed by different pipelines, the total-column score (TC-score), which was first introduced in BAliBASE [41], is the most popular measurement in many alignment benchmark tests. TC-score represents the percentage of the correctly aligned columns in alignments comparing with the references. Qscore (http://www.drive5.com/qscore) is an essential tool for analyzing the quality of MSAs in this paper. We chose the famous BAliBASE, OXBench, and SABmark benchmarks as the evaluation materials.

Table II summarizes for each pipeline 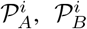 or 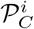 and each benchmark database the average TC-scores of the alignments constructed by the pipelines for the families in the database.

**TABLE II:**
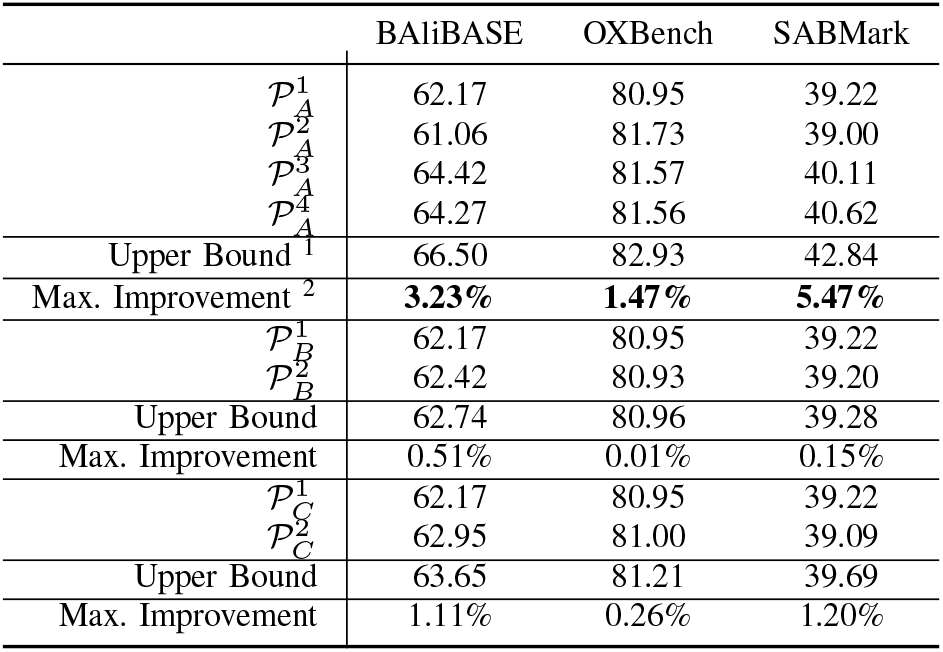
Average TC-score of each tool on the three empirical benchmarks

Table II shows that if a particular decision is used in Part A to assist in selecting different calculation methods, the theoretical maximum promotion proportion can be obtained. So next, we chose the right decision-making method for pipeline 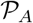.

### B. Determine the best decision-making model

We implemented the neural network structures mentioned in Section II-B2 and named them CNN, CNN + RNN, CNN + LSTM, CNN + BiLSTM, CNN + GRU and CNN + BiGRU, according to the different recurrent neural network layers used.

We divided the collected pairs data into two subsets: (1) 80% was randomly selected for model training, and (2) the remaining 20% was used for final testing. In the training process, a five-fold cross-validation was performed. This kind of validation method proved to be the most efficient [42]. In the process of training, we also set early stopping to further reduce overfitting.

Table III reveals the macro average (averaging the unweighted mean per label) [43] of the precision, recall and f_1_-score of the four labels in the 20% test data. If 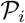 is the precision and 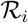 is the recall rate of class *i*, where *i* = 1, 2, 3 or 4 in this paper, the macro average precision 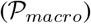 and recall 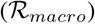 can be calculated by Formula (3).

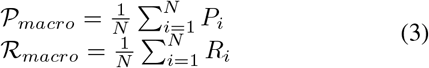

where *N* = 4, which means the category number is 4 in this paper.

**TABLE III:** The macro average precision, recall and f_1_-score on the test data

| Models | $\mathcal{P}_{macro}$ | $\mathcal{R}_{macro}$ | $\mathcal{F}_{macro}$ |
| --- | --- | --- | --- |
| CNN | 87.13 | 86.69 | 86.91 |
| CNN + RNN | 86.90 | 88.13 | 87.51 |
| CNN + LSTM | 87.13 | 87.71 | 87.41 |
| **CNN + BiLSTM** | 87.70 | 88.68 | **88.19** |
| CNN + GRU | 87.93 | 86.71 | 87.32 |
| CNN + BiGRU | 88.23 | 87.89 | 88.06 |

The macro average f_1_-score 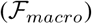 equals to the harmonic average of 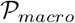 and 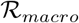.

According to the f_1_-score in Table III, we decided to use CNN + BiLSTM as our decision-making model.

Table IV illustrates, in more detail, the precision, recall and f_1_-score in the four different categories of the CNN + BiLSTM model we finally selected.

**TABLE IV:** The precision, recall and f_1_-score on the test data in different categories

| Category | Precision | Recall | $F_1$-score |
| --- | --- | --- | --- |
| $\mathcal{P}_A^1$ | 82.97 | 96.90 | 89.39 |
| $\mathcal{P}_A^2$ | 79.46 | 92.83 | 85.62 |
| $\mathcal{P}_A^3$ | 95.44 | 77.81 | 85.73 |
| $\mathcal{P}_A^4$ | 92.93 | 87.18 | 89.96 |
| Macro Avg. | 87.70 | 88.68 | 88.19 |

Although there is a difference in the precision or recall rate among the different categories, the f_1_-scores of each category is generally good; they were all over 85%, indicating that our model can handle all categories well.

Figure 2 further shows the confusion matrix obtained using CNN + BiLSTM as the decision-making model to predict the four categories of pipeline 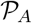. The confusion matrix is a table that is often used to describe the performance of a classifier on a set of test data for which the true values are known.

**Fig. 2:**
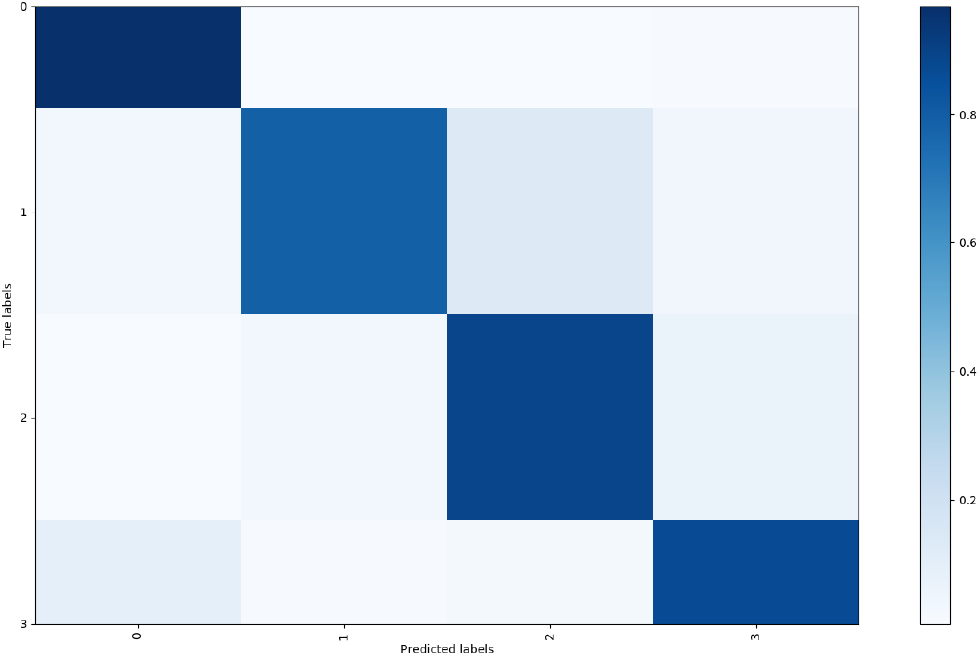
The confusion matrix of the decision-making model trained by CNN + BiLSTM. *The darker the color of the grid of the predicted label and the corresponding true label, the higher the accuracy of the prediction of this category (category 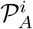 is represented by the label i* – 1).

The structure of the model we used is shown in Figure 3.

**Fig. 3:**
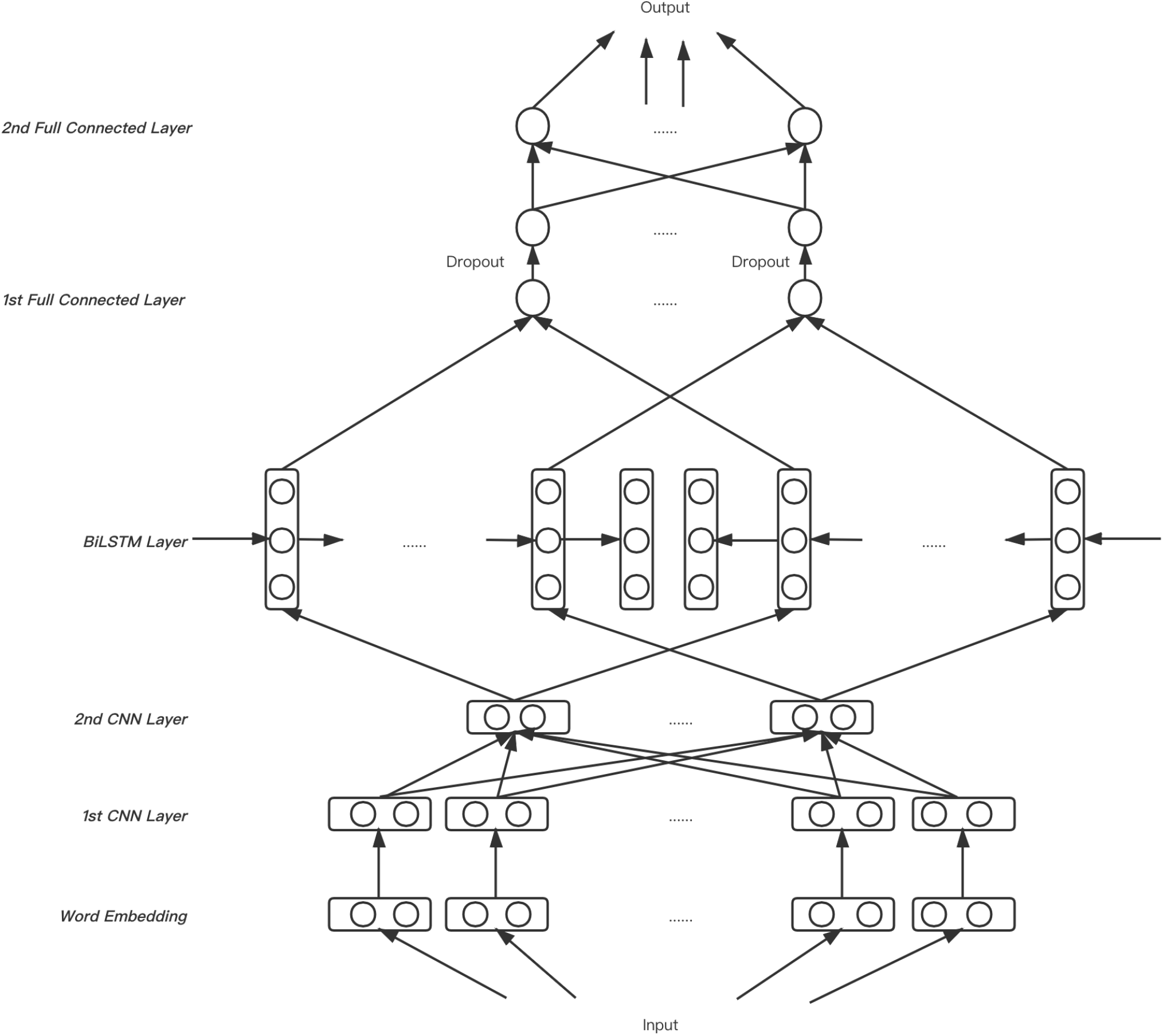
The neural network structure of our decision-making model. *Firstly the input is transformed through the Word embedding layer into a 512* × *8 matrix. Then the matrix passes through two CNN layers with filter sizes of 6 and 3 respectively (each CNN layer is followed by a max-pooling layer of size 2). Next, the output of the previous layer goes through the Bi-directional LSTM layer with a hidden size of 64. Finally, two full connection layers are connected and a 0.5 dropout to the first full connection layer is set.*

### C. DLPAlign: Decision-making model-based progressive alignment method

Given the high accuracy of our decision-making model (CNN + BiLSTM above), we integrated it into existing progressive alignment methods to construct a new alignment tool for multiple protein sequences, which we named DLPAlign. Given a protein family 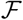, DLPAlign produces a multiple sequence alignment using mainly Steps III-C1 - III-C5.

#### 1) Decision-making for the posterior probability matrix calculation

Each pair *x, y* in 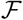 is inserted into the model with the highest accuracy described in the previous section to get a label (say *label_x;y_*). These labels represent the specific calculation method of posterior probability matrix that should be used. Which calculation method is chosen for the protein family 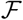 is determined by the dominate proportion of labels that all of its pairs get after passing through the decisionmaking model.

Because we already know the percentage of each correct label of 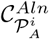 where *i* = 1, 2, 3 or 4, the dominate proportion of predicted labels can be calculated using the following Formula (4).

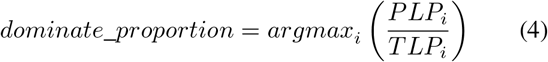

where *PLP_i_* means the percentage of the *i^th^* predicted label, *TLP_i_* means the proportion of the *i^th^* true label and *i* = 1, 2, 3 or 4.

This process is shown in Figure 4.

**Fig. 4:**
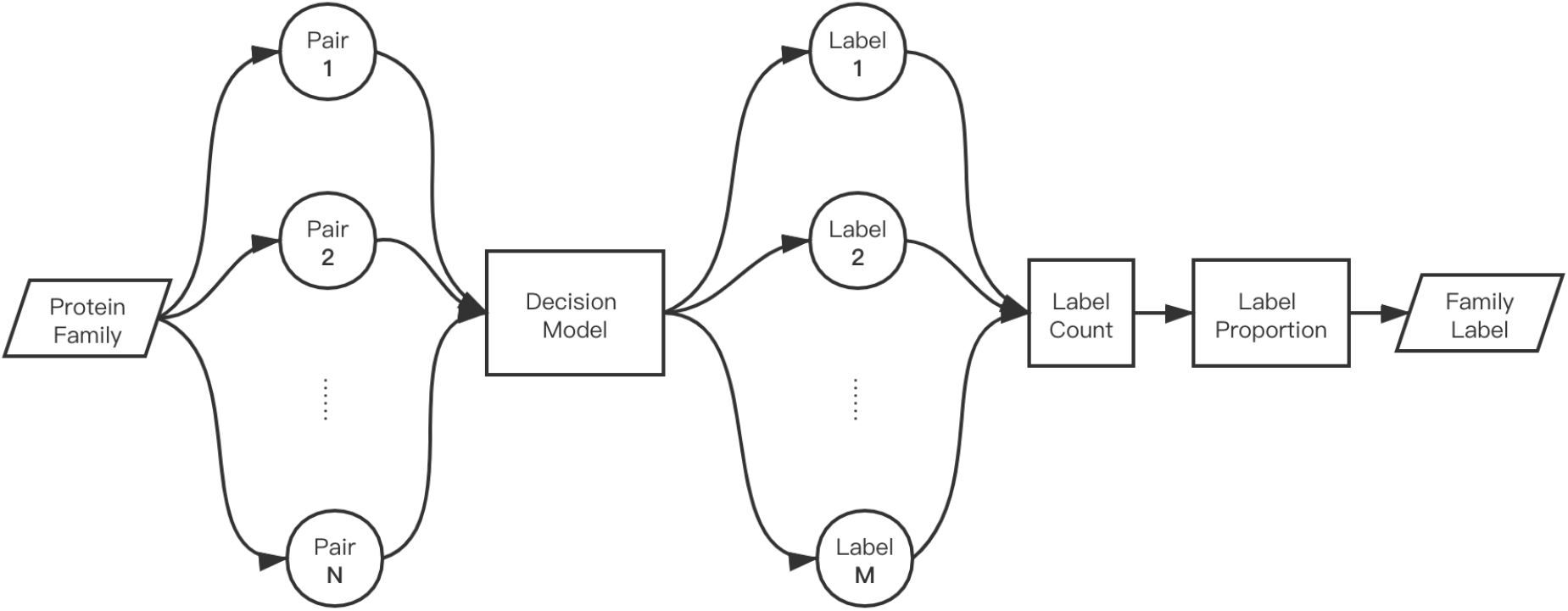
The process of splitting a protein family into pairs, using decision-making model to determine the label of each pair, and finally calculating the dominate proportion of the labels to get the label of the protein family.

Depending on the final family category, we use (1) the pair HMM, (2) the partition function, (3) the RMS of pair HMM and the partition function or (4) the RMS of pair HMM,the partition function and random HMM to accomplish the calculation of the posterior probability matrix.

#### 2) Distance matrix calculation and guide tree generation

By finding the maximum summing path (or maximum weight trace) through the posterior probability matrix, a pairwise alignment is computed on every pair *x*, *y* in 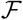 and the maximum sum is saved as *Prob*(*x, y*). The distance between sequences *x* and *y* can be measured by the Equation (5).

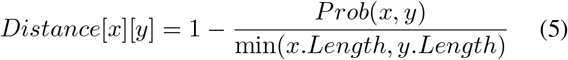

where *x.Length* and *y.Length* represent the length of sequences *x* and *y*, respectively.

A guide tree is a data structure used to determine the relationship between (1) sequence and sequence, (2) sequence and profile, and (3) profile and profile. Defining two clusters A and B, the distance between their union and another cluster C can be expressed as Formula (6), which is also the specific implementation of UPGMA.

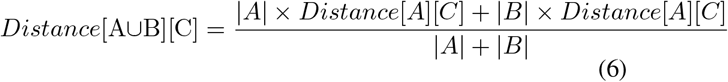

where |*A*|, |*B*| and |*C*| represent the weight of clusters A, B and C.

According to this distance calculation formula and the distance matrix computed, we can start from the sequences of the minimum distance and gradually builds a binary tree, named “Guide Tree”.

#### 3) Consistency transformation

In this step, we use other sequences to relax the posterior probability matrix of every pair *x* and *y* (written as *P_x,y_*), which we calculated in Step III-C1 to determine the substitution scores for the relaxation. The relaxation process can be expressed by the Equation (7).

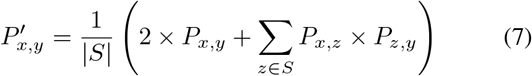

where *S* stands for the sequences set in protein family 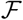, and 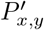 is the new transformed posterior matrix of pair < *x,y* >.

#### 4) Progressive alignment

Based on the guide tree we determined in the Step III-C2 and the relaxed posterior probability matrix in the Step III-C3, we can merge two child nodes (sequences) from the deepest node to get a profile, and then merges them to the root node of the tree respectively to get a complete MSA containing all sequences in 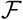.

#### 5) Refinement

The purpose of refinement is to correct some errors that may have occurred in the alignment between previous sequences. In this specific implementation, we also used the iterative refinement step to divide all aligned sequences into two groups each time randomly and then used the profile-profile alignment to realign them again. However, we added accuracy judgment. Each refinement was valid only if the maximum sum described in Section III-C2 was larger than before.

## IV. Results

### A. Comparing the accuracy of DLPAlign with other MSA tools

To determine the accuracy of DLPAlign implemented by the CNN + BiLSTM decision-making model and comparing this with other MSA tools with high accuracy, three empirical benchmarks were selected - BAliBASE 3.0, OXBench 1.3 and SABmark 1.65 - and the newest versions of eleven popular MSA tools were chosen for comparison: QuickProbs, PnpProbs, GLProbs, MSAProbs, Probalign, ProbCons, PicXAA, MAFFT, MUSCLE, Clustal Ω and T-Coffee. Of these eleven MSA tools, PicXAA adopted the non-progressive strategy, PnpProbs used both the non-progressive strategy and the progressive strategy, and the others used the progressive strategy. The TC-score mentioned in Section III-A was the leading indicator in the comparison.

The alignments in BAliBASE were organized into reference sets that were designed to represent real multiple alignment problems. Table V shows the average TC-score of the whole benchmark with 386 families and the accuracy of its two divergent reference sets (say, RV11 and RV12). DLPAlign could also handle RV11, which is a very divergent subset, obtaining a 1.58% higher TC-score than the second-best MSA tool.

**TABLE V:** Average TC-scores for BAliBASE

|  | All (386) | RV11 (38) | RV12 (44) |
| --- | --- | --- | --- |
| DLPAlign | **65.47** | **47.67** | **87.32** |
| QuickProbs | **65.41** | **46.93** | 87.03 |
| PnpProbs | 62.46 | 45.15 | 87.25 |
| GLProbs | 62.09 | 44.68 | **87.38** |
| MSAProbs | 64.51 | 44.40 | 87.03 |
| ProbCons | 61.89 | 40.89 | 84.14 |
| PicXAA | 59.97 | 46.64 | 86.60 |
| MAFFT | 50.08 | 28.23 | 75.57 |
| Muscle | 53.17 | 32.06 | 58.90 |
| ClustalΩ | 56.20 | 36.22 | 79.38 |
| ProbAlign | 60.68 | 45.69 | 86.69 |
| T-Coffee | 59.16 | 39.27 | 84.16 |
| Improved % | 0.09 | 1.58 | -0.07 |

Table VI shows the results of DLPAlign, as well as other MSA tools, for OXBench. In addition to the complete set of families, the table shows the alignment accuracy for families with average PID of less than and more than 30%.

**TABLE VI:** Average TC-scores for OXBench

|  | All (395) | 0 - 30% (63) | 30% - 100% (332) |
| --- | --- | --- | --- |
| DLPAlign | **82.52** | **44.16** | **89.80** |
| QuickProbs | 81.77 | 41.50 | 88.35 |
| PnpProbs | **82.06** | **43.96** | 89.54 |
| GLProbs | 81.93 | 43.34 | **89.55** |
| MSAProbs | 81.50 | 42.81 | 89.08 |
| ProbCons | 80.68 | 41.50 | 88.35 |
| PicXAA | 81.14 | 39.78 | 89.32 |
| MAFFT | 78.15 | 35.93 | 86.40 |
| Muscle | 80.67 | 40.95 | 88.21 |
| ClustalΩ | 79.99 | 37.39 | 88.08 |
| ProbAlign | 81.68 | 41.06 | 89.39 |
| T-Coffee | 80.18 | 40.04 | 87.80 |
| Improved % | 0.56 | 0.45 | 0.28 |

Note that OXBench did not divide the whole database into different subsets. We made the division here because we thought the two parts after separation could respectively represent divergent families and high similarity families.

It can be seen that no matter which divergent set or high similarity set is used, DLPAlign can always produce some improvement, although the improvements are sometimes small.

Table VII summaries the accuracy for SABMark benchmark, which was divided into two subset Twilight Zone and Superfamilies, depending on the SCOP classification. These subsets together covered the entire known fold space using sequences with very low to low, and low to intermediate similarity, respectively. DLPAlign improved both subsets and the whole benchmark. For Twilight Zone, except for DLPAlign, none of the MSA tools could get a TC-score of more than 25%. In this subset, DLPAlign’s TC-score was 4.38% higher than the second-best MSA tool, PnpProbs. The results were very surprising.

**TABLE VII:** Average TC-scores for SABMark

|  | All (423) | Superfamily (315) | Twilight Zone (108) |
| --- | --- | --- | --- |
| DLPAlign | **42.59** | **48.37** | **25.71** |
| QuickProbs | 40.65 | 46.56 | 23.40 |
| PnpProbs | **41.48** | **47.03** | **24.63** |
| GLProbs | 41.22 | 46.95 | 23.86 |
| MSAProbs | 40.04 | 45.81 | 22.60 |
| ProbCons | 39.17 | 44.64 | 22.57 |
| PicXAA | 38.44 | 44.45 | 20.33 |
| MAFFT | 33.00 | 39.10 | 14.72 |
| Muscle | 33.47 | 39.13 | 16.96 |
| ClustalΩ | 35.47 | 41.43 | 18.10 |
| ProbAlign | 38.63 | 44.11 | 22.64 |
| T-Coffee | 39.07 | 44.15 | 24.25 |
| Improved % | 2.68 | 2.85 | 4.38 |

### B. Efficiency results

All the tools were run on an HP desktop computer with four Intel Cores i5-3570 (3.40 GHz) and a main memory of size 19.4 GB. Table VIII shows the efficiency results.

**TABLE VIII:** Average running time (in seconds) of three benchmarks by DLPAlign and other MSA tools

|  | Average running time (sec.) |  |  |
| --- | --- | --- | --- |
|  | BALiBASE | OXBench | SABMark |
| DLPAlign | 38.10 | 0.84 | 0.40 |
| QuickProbs | 6.10 | 0.11 | 0.06 |
| PnpProbs | 13.30 | 0.27 | 0.21 |
| GLProbs | 13.63 | 0.23 | 0.15 |
| MSAProbs | 9.49 | 0.14 | 0.07 |
| ProbCons | 33.98 | 0.49 | 0.24 |
| PicXAA | 29.82 | 0.49 | 0.35 |
| MAFFT | 0.33 | 0.15 | 0.14 |
| Muscle | 1.69 | 0.04 | 0.08 |
| ClustalΩ | 0.97 | 0.03 | 0.04 |
| ProbAlign | 21.92 | 0.27 | 0.12 |
| T-Coffee | 339.6355 | 6.1013 | 2.8393 |

It should be noted that ProbCons, Probalign, MSAProbs, GLProbs and PnpProbs all used the standard progressive alignment steps mentioned in Section **??**, so there was not much difference in their running times. The running time of DLPAlign comprised mainly the running time of the decisionmaking model and the standard progressive alignment. As Table VIII shows, the time taken by the decision-making model was affected by the benchmark size (BAliBASE was largest benchmark while SABMark was the smallest), but in general, the time was acceptable.

### C. A real-life application: Protein Secondary Structure Prediction

Protein secondary structure prediction is an appropriate application of multiple sequence alignment [44].

We picked some protein sequences related to SARS-COV-2 which could be found at Protein Data Bank (PDB) [45] with these PDB ID: 6YI3, 6VYO, 6W9C and 6W61 ^1^. We then used the following process to evaluate the performance of DLPAlign with the other five highly accurate MSA tools.

> Given a protein sequence *S*, we use Jpred 4 [46] to search protein sequences similar to it. Then, we construct an MSA 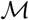 for these sequences and *S*, and use the secondary structure prediction tool provided on Jpred 4 to predict the secondary structure of *S*. Finally, comparing the predictions with reference secondary structures offered by Jpred 4.
>
> Table IX summaries the number of wrongly aligned residues for each MSA tool. The table shows that DLPAlign always got the fewest wrong aligned number of the leading MSA tools.

**TABLE IX:** Number of wrongly aligned residues in the predicted secondary structures of proteins

| PDB ID (Length) | DLPAlign | QuickProbs | PnpProbs | GLProbs | MSAProbs | PicXAA |
| --- | --- | --- | --- | --- | --- | --- |
| **6YI3 (140)** | **10** | 16 | 12 | 12 | 15 | 16 |
| **6VYO (128)** | **5** | **5** | **5** | **5** | **5** | 7 |
| **6W9C (317)** | **20** | 31 | 24 | 22 | 23 | 32 |
| **6W61 (299)** | **13** | 17 | **13** | **13** | 15 | 20 |

## V. Conclusion

The significant contributions of this paper are (i) using deeplearning methods to train a decision-making model to determine which specific calculation method to use in the posterior probability matrix calculation of progressive alignment approaches and (ii) releasing a new progressive multiple protein sequence alignment tool based on this deep learning model named DLPAlign. As hindsight after our study, DLPAlign with decision-making model could get better accuracy on all tests and perform especially excellent on low similarity families.

We would also like to point out that we have not optimized DLPAlign for efficiency. The efficiency of DLPAlign slows down with the increase of the number of sequences, because the input of the decision-making model is sequence pair. The more sequences, the more sequence pairs will be obtained; thus, the number of times the decision-making model runs is increased.

One way to improve efficiency is to reduce the number of times the decision-making model runs. In this study, we regard each sequence pair as a sentence to classify. If we can take the combination of more than 2 sequences as the input of the decision-making model, then the whole decisionmaking model will run less, which will significantly improve the efficiency of DLPAlign. It is also one of the improvements of DLPAlign in the future.

Besides, if we use the protein family simulation tool such as INDELible [47] to obtain enough protein family data, we can also consider the entire protein family or the temporary MSA built by the fast MSA tool as the training data of the decisionmaking model, so that the decision-making model will only run once, the efficiency can be further improved.

## VI. Acknowledgment

This research was guided by Dr. Hing-fung Ting of the University of Hong Kong. We thank Dr. Ting and Dr. Bin Yan from the University of Hong Kong for many helpful discussions and suggestions.

## Footnotes

1 Upper Bound: The highest TC-score could be obtained when each family chooses the pipeline which could get the best TC-score in this part.

2 Max. Improvement: The proportion that the upper bound can improve compared with the best-existed result of this part.

1 They are all available only from April 2020, and relevant references have not been published.

